# Signed, Sealed, Delivered: A Generalizable Model for Probiotic Delivery and Metabolism in the Gut

**DOI:** 10.1101/2025.06.20.660673

**Authors:** Vicenzo L. DeVito, Bhargav R. Karamched

## Abstract

As genetic engineering advances, so does the need for quantitative models to inform biological engineering. This is particularly true for the nascent field of synthetic probiotic therapy of the gut microbiome. Gut microbiome health is linked to health of the host organism. Decline in microbiome health is correlated with severe metabolic disorders. However, as far as we know, there are no models that account for ingestion and transfer of a synthetic probiotic as well as its intended metabolic effect. We present here the first model that accounts for such effects. We call our model the bacterial compartment absorption and transit (BCAT) model. It is a generalization of the pharmacokinetic compartment absorption and transit (CAT) model As a specific example, we employ the BCAT model in the context of trimethylaminuria (TMAU), a metabolic disorder characterized by a persistent fishy odor emanating from affected individuals. The BCAT model predicts the dose of probiotic required for adequate medical treatment of TMAU. Moreover, our model is flexible to apply to any metabolic disorder pertaining to the gut microbiome.

## 1 Introduction

With significant advancements in genetic engineering and synthetic biology over the past twenty years, development of health technology has grown tremendously. Synthetic microbial consortia have, in particular, shown promise as delivery mechanisms for biomolecular therapy [1–6]. Engineered bacteria such as *E. coli* [7–9], *L. lactis* [10–16], and *B. ovatus* [10, 17] have been introduced to the gut microbiome as synthetic probiotics as a means of silencing oncogenes, inhibiting proteases, and killing harmful pathogens such as *Pseudomonas aeruginose* [18]. Synthetic probiotics (hereafter used interchangeably with ‘probiotics’) are also being used as metabolic therapeutics, aiding expression of key biomolecules that help mitigate metabolic disorders such as diabetes [19–21]. This is particularly significant because a large percentage of metabolism incepts in the gut microbiome, which has a metabolic capacity—containing over three million genes—larger than that of the liver [18, 22–25].

As synthetic probiotics develop and become mainstream in biomolecular therapy, quantitative tools that inform and optimize therapeutics will become increasingly important for them. Indeed, integration of experimental and theoretical techniques has been successful in accelerating understanding of, for example, the role biomolecular oscillations play in coordinating activity across engineered microbial consortia, especially in spatially extended domains [2, 26–30]. Mathematical modeling has not been as integrated in biomolecular therapy of the gut microbiome. In particular, as far as we are aware, there has been no mathematical model developed that models transfer of probiotic through the gastrointestinal (GI) tract as it performs a desired metabolic function.

In this short paper, we present the first mathematical model of probiotic metabolism in conjunction with its transfer through the small intestine. Our model is based on the compartment absorption and transfer (CAT) model prevalent in physiologically-based pharmacokinetic (PBPK) modeling [31–33]. The CAT model effectively treats the small intestine as a 1D spatial domain and tracks the dynamics of ingested food passing through. The boundaries for the domain occur at the stomach and the colon. The CAT model functionally acts as a discretization of an advection-degradation partial differential equation. If *ρ*(*x, t*) describes the concentration of a substance at location *x* ∈ (0*, L*) of the digestive tract at time *t >* 0, then the dynamics of the substance obey

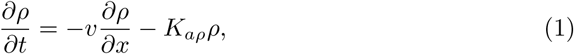

where *v* describes the velocity of transport through the digestive tract and *K_aρ_* is the rate of absorption of substance *ρ* across the lining of the small intestine into the plasma. Replacing the spatial derivative with an upwind discretization [34, 35] yields

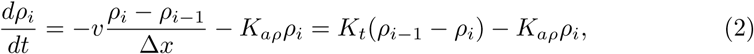

where now *ρ_i_*(*t*) is the concentration of a substance in the *i*th compartment, with 1 ≤ *i* ≤ *N* for some *N* ∈ N, and Δ*x* is the fundamental length unit used to define the compartments. The quantity *K_t_* ≡ *v/*Δ*x* is the inverse of the residence time of the substance within each compartment. Equation (2) is coupled with the following equations for the concentration of the substance in the stomach, *ρ*_s_,the colon, *ρ*_c_, which describe the boundary dynamics, and the plasma *ρ*_pl_, where substances go after absorption through the intestinal wall:

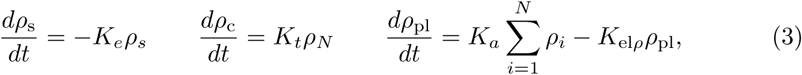

where *K_e_* is the gastric emptying rate constant and *K*_el_*_ρ_* represents degradation of substance *ρ*. Equations (2)–(3) together form the CAT model.

The CAT model is mathematically simple. It assumes that propagation of nutrients through the digestive tract occurs at constant speed and that absorption of nutrients across the membrane of the small intestine is homogeneous. Furthermore, using *N* = 7 compartments was found to be the optimal formulation of the CAT model [36–40]. Despite its mathematical simplicity, the CAT model has proved to be a successful pharmacokinetic model. That is, it has been used to predict whether or not a specific dose of an intervention suffices to achieve a desired concentration in the plasma. Furthermore, it has been the foundation for other successful pharmacokinetic models, such as the ACAT model [41], which is the industry standard for softwares such as GastroPlusTM [42].

We introduce a simple modification to the CAT model by explicitly incorporating dynamics of probiotic, a given metabolite, the corresponding metabolism, and the impact of natural gut bacteria upon the metabolism. We call our new model the bacterial CAT model (BCAT). In our model, probiotic expresses an enzyme that metabolizes the metabolite under scrutiny, but we do not incorporate enzyme dynamics explicitly. We assume that enzyme is at equilibrium and employ Michaelis-Menten kinetics to describe metabolism. Our model predicts dosage of probiotic required to meet benchmarks of a specific intervention.

The BCAT model is mathematically simple and flexible so that dynamics of any engineered probiotic and the corresponding metabolic activity can be incorporated into its framework. Estimation of parameters pertaining to absorption and transit through the small intestine are highly uncertain. But the computational efficiency of the BCAT model allows for optimized parameter sweeps that facilitate parameter selection so that model outputs are biologically plausible.

We will demonstrate two fundamental features of the BCAT model in this paper: (1) its ability to predict dosage of an intervention required to attain specific desired biomedical metrics and (2) its ability to predict optimal timing of dosage administration.

To demonstrate the utility of the BCAT model, we specifically employ it in the context of trimethylaminuria. Trimethylaminuria (TMAU) is a rare metabolic disorder characterized by a persistent fishy odor emanating from affected individuals [43]. This condition arises from mutations in the FMO3 gene, which encodes flavin-containing monooxygenase 3 (FMO3)—an enzyme essential for converting odorous trimethylamine (TMA) into its non-odorous oxidized form, trimethylamine *N* -oxide (TMAO) [44]. Deficiencies in the FMO3 enzyme lead to the accumulation of TMA in the bloodstream, which is eventually excreted in bodily fluids, causing the characteristic odor. TMA can enter the bloodstream through two primary pathways: (1) direct consumption of foods rich in TMA, such as seafood [45]; or (2) consumption of precursors like choline, betaine, and L-carnitine, which gut microbiota metabolize into TMA [46, 47]. TMA is subsequently absorbed through the intestinal wall and transported to the liver via the portal vein.

Epidemiological data estimate that TMAU affects between 1 in 40,000 and 1 in 1,000,000 individuals worldwide [48, 49]. While the condition is not life-threatening, its impact on quality of life is profound. Individuals with TMAU often face social isolation, difficulties in maintaining employment, and strained interpersonal relationships due to the symptoms and stigma associated with their condition. Psychological consequences, including low self-esteem, depression, and anxiety, are common [50].

Currently, there are no definitive cures or universally effective treatments for TMAU. Common management strategies involve lifestyle changes, such as avoiding foods high in TMA and choline and ingesting charcoal and copper supplements, or post-hoc treatments, such pH-balanced soaps to neutralize scent or antibiotics to reduce TMA-producing gut microbes [43, 44, 51]. Such treatment strategies do not address the underlying issue and have acute downsides. Avoiding choline-rich foods is a plausible preventative intervention, but the modern diet is abundant with choline-rich foods [52, 53]. Such a diet alteration would be drastic. On the other hand, taking antibiotics to prevent gut microbiota metabolism of choline into TMA is suboptimal due to the loss of natural gut bacteria essential for metabolic health [18]. Novel therapeutics are therefore required for sustainable TMAU management.

Indeed, we initially set out to develop a probiotic treatment for TMAU with genetically modified bacteria. To determine appropriate dosage for such a treatment, we scanned the literature for mathematical models that predict optimal dosage. To our surprise, no such model was available. Specifically, to our knowledge and through our findings, we could not find a mathematical model that accounted for both the pharmacokinetics and the pharmacodynamics of a probiotic intervention. This drove our development of BCAT. As far as we know, this is the first model to describe natural gut bacterial effects as well as probiotic dynamics in the context of metabolism—specifically a metabolic disorder.

We propose to have genetically-engineered probiotics express trimethylamine monooxygenase (TMM)—an enzyme which is effectively the bacterial version of FMO3 in that it oxidizes TMA into TMAO [54]. We hypothesize that ingesting a dose of such probiotic with a meal allows for TMA oxidation in the digestive tract prior to transfer to the liver via the portal vein. We use the BCAT model to predict an exact dosage of probiotic required to achieve TMAO levels that are comparable to people without TMAU. Thus, the BCAT model is a hybrid pharmacokinetic/pharmacodynamic model, capable of predicting whether or not a given dosage of intervention yields desired plasma concentration levels and conversely predicting dosages to achieve a desired output. Our model’s dosage prediction is consistent with what can be achieved in common settings.

## 2 The BCAT Model

We now present the BCAT model. Because choline is a major player in consumption of TMA in the modern diet, we include choline in our model of TMA metabolism. This necessitates the inclusion of intestinal bacteria in our model as well because gut microbes metabolize choline into TMA. Let *c_i_*(*t*), *ϕ_i_*(*t*), and *ψ_i_*(*t*) denote the concentrations of choline, TMA, and TMAO in the *i*th compartment at time *t*, respectively. Let *b_i_* and *p_i_*(*t*) denote the concentrations of intestinal bacteria and probiotics in the *i*th compartment at time *t*, respectively. Variables with subscripts ‘s’, ‘co’, and ‘pl’ denote the corresponding concentrations in the stomach, colon, and plasma, respectively. The full model can be seen in Table 1.

**Table 1:**
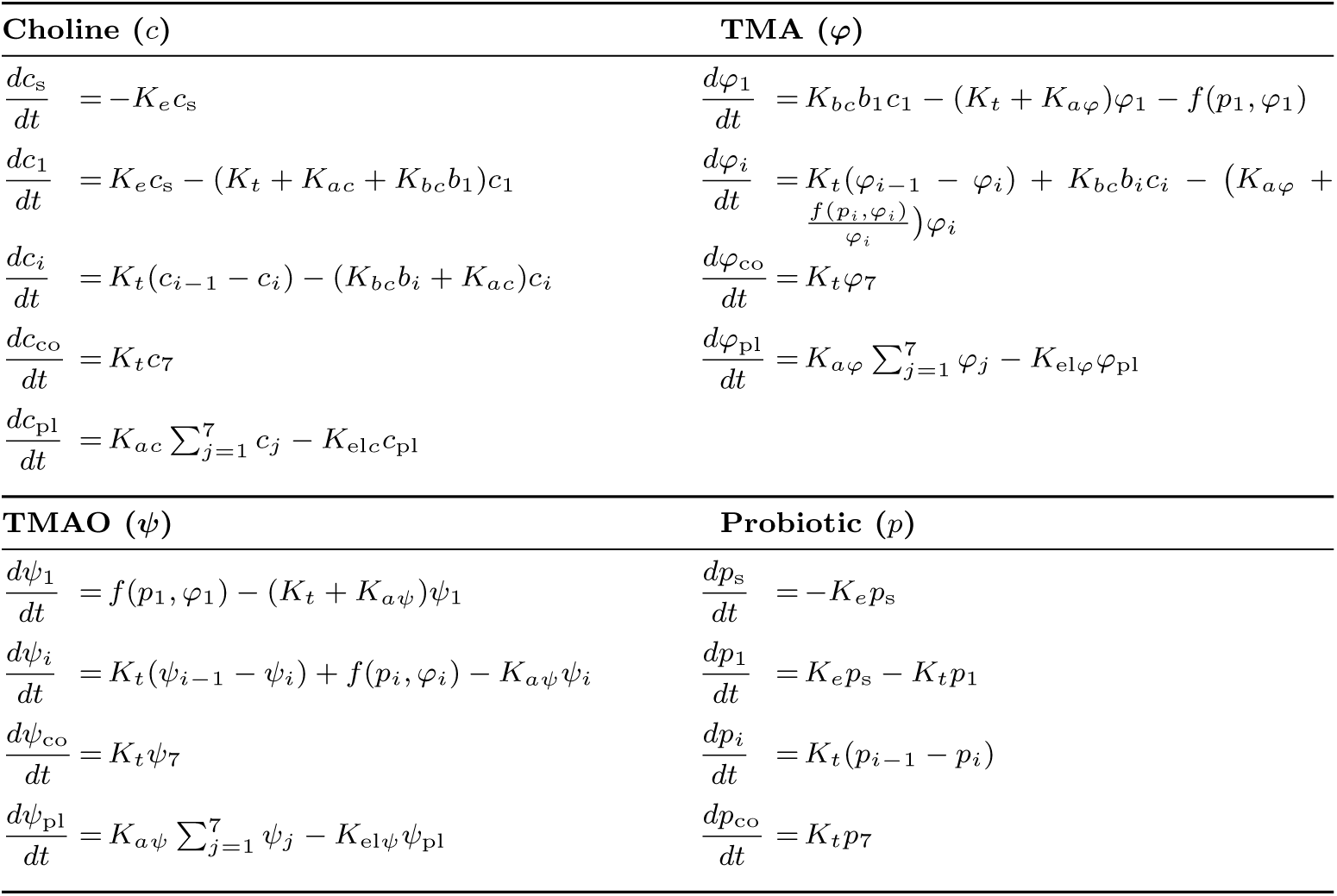
BCAT model system of ordinary differential equations.

| **Choline ($c$)** | **TMA ($\varphi$)** |
| --- | --- |
| $\frac{dc_s}{dt} = -K_e c_s$ | $\frac{d\varphi_1}{dt} = K_{bc} b_1 c_1 - (K_t + K_{a\varphi}) \varphi_1 - f(p_1, \varphi_1)$ |
| $\frac{dc_1}{dt} = K_e c_s - (K_t + K_{ac} + K_{bc} b_1) c_1$ | $\frac{d\varphi_i}{dt} = K_t (\varphi_{i-1} - \varphi_i) + K_{bc} b_i c_i - (K_{a\varphi} + \frac{f(p_i, \varphi_i)}{\varphi_i}) \varphi_i$ |
| $\frac{dc_i}{dt} = K_t (c_{i-1} - c_i) - (K_{bc} b_i + K_{ac}) c_i$ | $\frac{d\varphi_{co}}{dt} = K_t \varphi_7$ |
| $\frac{dc_{co}}{dt} = K_t c_7$ | $\frac{d\varphi_{pl}}{dt} = K_{a\varphi} \sum_{j=1}^7 \varphi_j - K_{el\varphi} \varphi_{pl}$ |
| $\frac{dc_{pl}}{dt} = K_{ac} \sum_{j=1}^7 c_j - K_{elc} c_{pl}$ | |
| **TMAO ($\psi$)** | **Probiotic ($p$)** |
| $\frac{d\psi_1}{dt} = f(p_1, \varphi_1) - (K_t + K_{a\psi}) \psi_1$ | $\frac{dp_s}{dt} = -K_e p_s$ |
| $\frac{d\psi_i}{dt} = K_t (\psi_{i-1} - \psi_i) + f(p_i, \varphi_i) - K_{a\psi} \psi_i$ | $\frac{dp_1}{dt} = K_e p_s - K_t p_1$ |
| $\frac{d\psi_{co}}{dt} = K_t \psi_7$ | $\frac{dp_i}{dt} = K_t (p_{i-1} - p_i)$ |
| $\frac{d\psi_{pl}}{dt} = K_{a\psi} \sum_{j=1}^7 \psi_j - K_{el\psi} \psi_{pl}$ | $\frac{dp_{co}}{dt} = K_t p_7$ |

The principal modifications of the CAT model yielding the BCAT is the inclusion of the bacterial and probiotic metabolic terms. We model the metabolism of choline by gut microbiota with the mass-action term while the enzymatic oxidation of TMA into TMAO is captured by the Michaelis-Menten term, where *V*_max_ is the maximum rate of oxidation, *E*_eq_ is the equilibrium concentration of TMM, and *K_m_* is the half-activation constant:

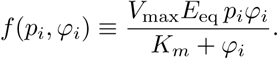

The model is fundamentally data-driven and was formulated around parameters and experimental observations reported in the literature; accordingly, probiotic activity is represented by Michaelis–Menten kinetics, whereas endogenous bacterial metabolism, due to the nature of reported relevant biophysical parameters in the literature, is captured with a mass-action term. A full list of parameters, their values, and literature sources is provided in Table 2

**Table 2:** Simulation parameters used in the BCAT model (rates in h^−1^)

| Parameter | Description | Value | Unit | Source |
| --- | --- | --- | --- | --- |
| $K_e$ | Gastric emptying rate | 0.852 00 | $\text{h}^{-1}$ | [59] |
| $K_t$ | Intestinal transit rate | 2.0690 | $\text{h}^{-1}$ | [33] |
| $K_{ac}$ | Absorption rate of choline | 2.8680 | $\text{h}^{-1}$ | [60] |
| $K_{a\varphi}$ | Absorption rate of TMA | 0.744 00 | $\text{h}^{-1}$ | [61] |
| $K_{a\psi}$ | Absorption rate of TMAO | 0.119 40 | $\text{h}^{-1}$ | [62] |
| $K_{elc}$ | Elimination rate of choline | 6.0000 | $\text{h}^{-1}$ | [63] |
| $K_{el\varphi}$ | Elimination rate of TMA | 0.300 30 | $\text{h}^{-1}$ | [64] |
| $K_{el\psi}$ | Elimination rate of TMAO | 0.173 00 | $\text{h}^{-1}$ | [65] |
| $K_{bc}$ | Bacterial conversion rate | $1.1000 \times 10^{-10}$ | $\text{ml CFU}^{-1} \text{h}^{-1}$ | [66] |
| $K_m$ | Michaelis–Menten constant | 0.021 600 | $\mu\text{mol}$ | [54] |
| $V_{\max}$ | Enzymatic reaction max. velocity | 68.016 | $\mu\text{mol mg}^{-1} \text{h}^{-1}$ | [54] |
| $E_{eq}$ | Equilibrium enzyme amount in bacteria | $1.0000 \times 10^{-9}$ | $\text{mg}$ | [67] |

We made the following simplifying assumptions to optimize computational efficiency of the BCAT model: (1) bacterial and probiotic populations throughout the full small intestine are constant; (2) TMA oxidation is instantaneous, meaning we ignore temporal delay arising from TMA diffusion across probiotic cell membrane; (3) TMM has already attained equilibrium in the probiotic cells, and thus its concentration within a probiotic cell is invariant. Assumption (1) means probiotic density through the compartments varies in time but Σ*_j_ p_j_* is constant, where *j* is taken over all compartments. The density of natural gut bacteria *b_i_* per compartment is taken to be invariant in time. We take *b_i_* = 10*^i^*^+1^ CFU/mL^1^ based on previously published reports [55]. Published reports on the values gut bacteria take are varied, and we find that *b_i_* can range between 10*^i^* and 10*^i^*^+2^ CFU/ml. Here, we will use the median in the model for simplicity.

We note that using standard Michaelis-Menten kinetics to describe enzymatic oxidation of TMA to TMAO is a modeling choice for this specific metabolic system and is completely generalizable for other more complicated enzymatic pathways. Incorporating competitive, noncompetitive, or even uncompetitive inhibitors to attain desired kinetics is easily captured by the BCAT model [56, 57].

Each state variable is prescribed an initial condition. All initial conditions are zero except *c*_s_(0) = *c*_0_ *>* 0 and *p*_s_(0) = *p*_0_ *>* 0. The initial data *p*_0_ is the dosage of the probiotic intervention prescribed. Unless otherwise specified, we take *c*_0_ = 10^3^*µ*mol in all simulations.

## 3 Simulation of Probiotic Intervention

We now employ the BCAT model to ascertain the optimal dosage *p*_0_. To define the optimal dosage, we consider the quantities

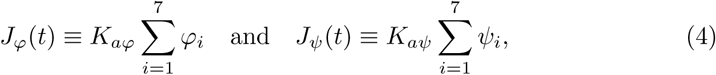

which quantitate the physicochemical fluxes of TMA and TMAO across the wall of the small intestine. The typical plasmal ratio of concentration of TMAO to total concentration of TMAO and TMA is 0.95. Thus, our objective is to determine the dosage *p*_0_ that gives

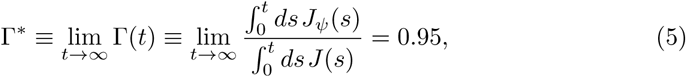

where

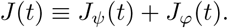

### 3.1 No treatment

In the baseline simulation, probiotic is not administered (*p*_0_ = 0), leading to the unrestrained conversion of choline into TMA and the subsequent absorption of TMA into the portal vein. In this case, Γ^∗^ ≡ 0. In Figure 2 we show time series of choline and TMA in the various compartments of the small intestine. The solutions resemble those of a linear transport equation with decay. That is, the solutions behave like a traveling wave with a decay constant determined by the absorption of the choline and TMA across the lining of the small intestine. Choline enters the small intestine from the stomach, so the temporal dynamics therein are given by an exponential decay of choline concentration. The dynamics throughout the remainder of the compartments

**Fig. 1:**
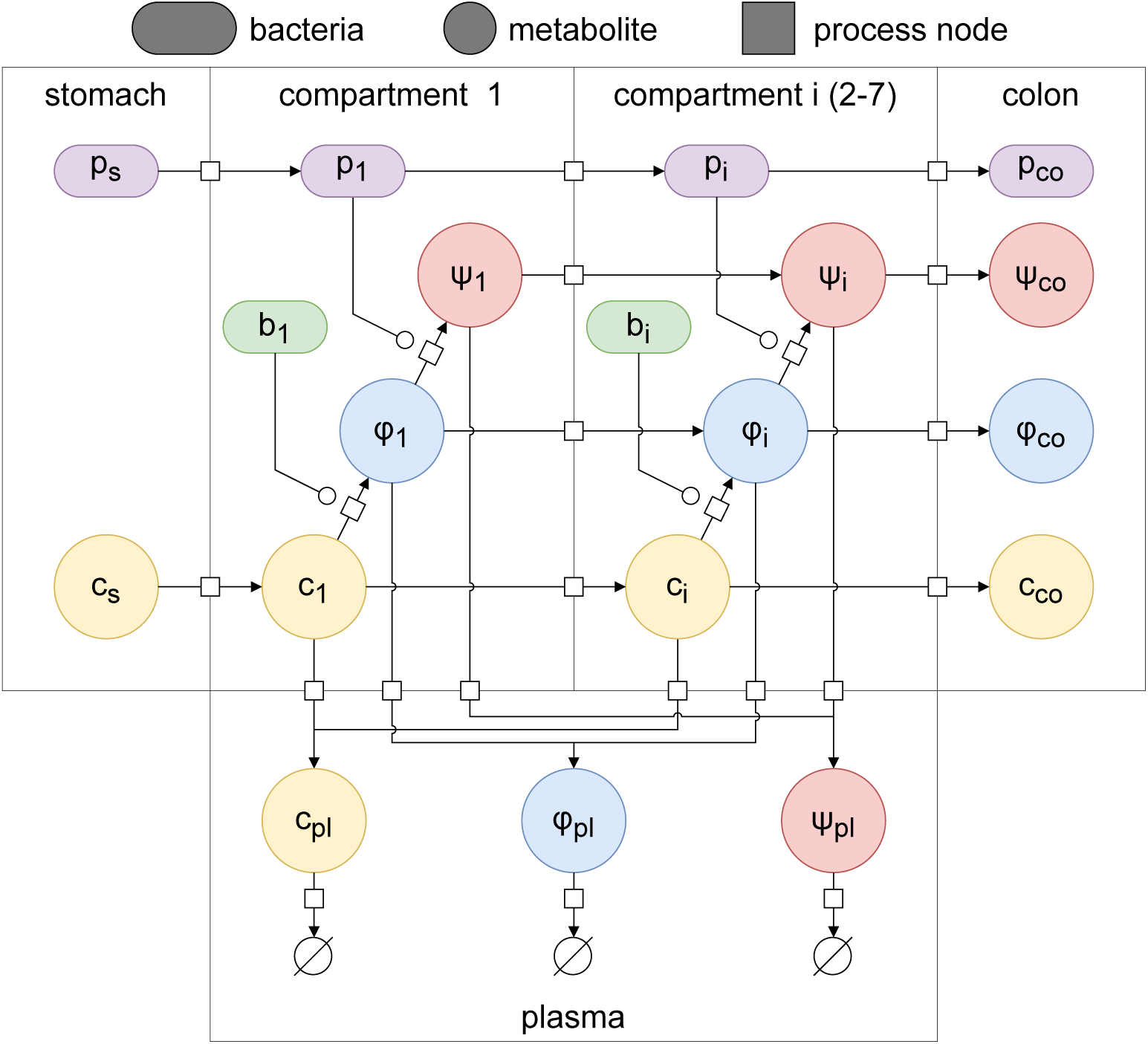
Schematic of the BCAT model. We use the Systems Biology Graphical Notation (SBGN) [58] to depict the interaction of the different species in the metabolic pathway.

**Fig. 2:**
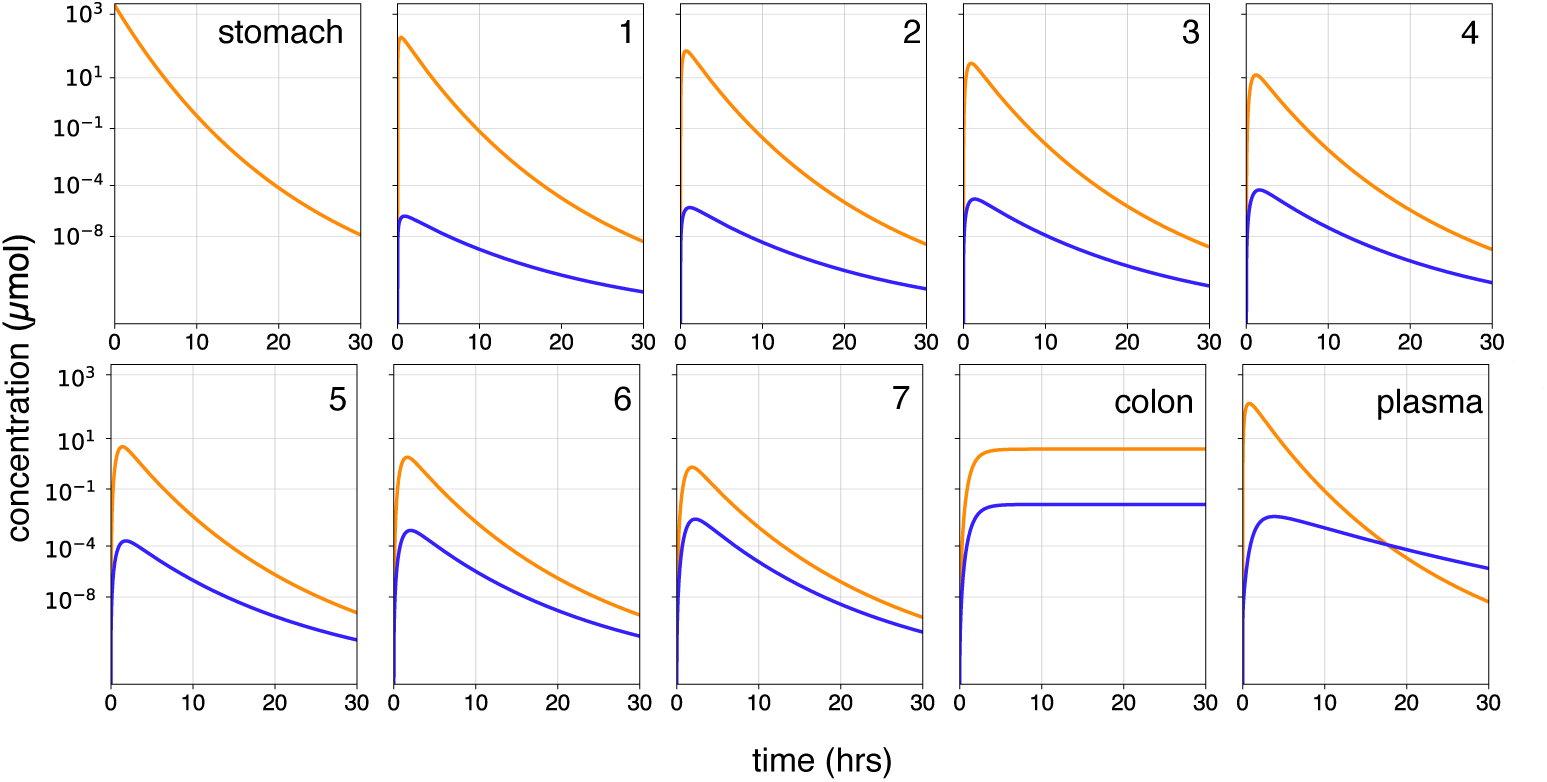
Time series of choline and TMA in each compartment according to the BCAT model (see Table 1) of the small intestine for the no treatment case. Parameter values are detailed in Table 2 and *c*_0_ = 10^3^ *µ*mol.

resemble a traveling wave solution with the dynamics of each compartment depicting a peaked time series wherein the choline concentration in compartment *i* tries to match the choline concentration in compartment *i* − 1 but fails due to choline absorption and metabolism. Dynamics in compartments 1-7 also show the emergence of TMA as a metabolic product of choline. The TMA dynamics also resemble a peaked response in all compartments except the colon, where we do not consider metabolic effects.

This leaves an important point of modeling in the future. While the role the large intestine plays in extracting nutrients from digested food is not quite as important as the small intestine, it can still affect interaction with ingested drugs, as indicated by the emerging field of pharmacomicrobiomics [68].

In the no treatment scenario, the BCAT model is linear and completely solvable.

Standard ODE theory tells us that

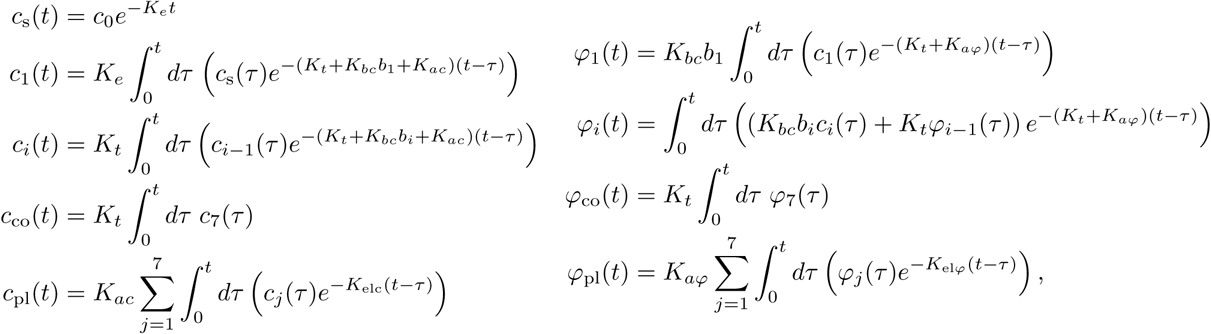

yielding the analytic result for TMA accumulation

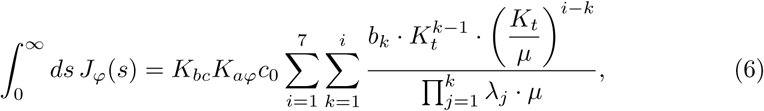

where *λ_j_* ≡ *K_ac_* + *K_t_* + *K_bc_b_j_* and *µ* ≡ *K_aϕ_* + *K_t_*. Though in this case we know TMA levels in the portal vein will be high, Eq. (6) unveils what biological parameters can control TMA absorption in the absence of probiotic metabolism. For example, decreasing the rate of absorption of TMA across the intestinal lining (*K_aϕ_* → 0) will clearly reduce TMA in the portal vein. Other ways to reduce TMA absorption in the absence of probiotic treatment as informed by Eq. (6) are (1) eliminate consumption of choline (*c*_0_ → 0), (2) prevent gut bacterial metabolism of choline into TMA (*K_bc_* → 0), (3) eliminate gut bacteria (*b_k_* → 0), (4) increase the choline rate of absorption across the small intestine lining (*K_ac_* → ∞), or (5) increase the rate at which TMA passes through the small intestine (*K_t_* → ∞, i.e., the residence time of TMA in each compartment converges to 0.). Indeed, currently available treatments for TMAU involve addressing these very parameters. People suffering from TMAU are encouraged to reduce choline consumption or to take antibiotics to reduce gut bacterial conversion of choline into TMA. However, as we discussed before, these treatments are unsustainable for successful management of TMAU. On the other hand, increasing the rate at which choline or TMA passes through the small intestine, or maximizing the absorption of choline across the GI lining are impractical treatments with no clear method of implementation.

### 3.2 Treatment

Here, we take *p*_0_ *>* 0 and generate a dose response curve to ascertain the minimum value of *p*_0_ yielding Γ^∗^ = 0.95. To generate the dose–response curve, we systematically increase the initial probiotic dose *p*_0_ and, for each value, solve the BCAT model to equilibrium. We then plot the resulting equilibrium ratio Γ^∗^ against *p*_0_. The minimal dose required to achieve Γ^∗^ = 0.95 is identified by locating the smallest *p*_0_ at which the curve intersects the line Γ^∗^ = 0.95. Importantly, we assume here that the probiotic is administered at the same time as the choline is ingested.

In Figure 3B-C, we show the impact of introducing probiotic on the physicochemical flux of TMA into the plasma. For dosage (*p*_0_) yielding Γ^∗^ = 0.95, we have approximately a two orders of magnitude less for peak TMA concentration in the plasma relative to the peak concentration in the absence of treatment. This decrease in TMA maps to a decrease in fishy odor emanating from affected individuals. In Figure 3D we show that the fraction of TMA to TMA and TMAO in the plasma stays bounded over time and quickly and inerratically attains its equilibrium value on the dose response curve. For *p*_0_ values

**Fig. 3:**
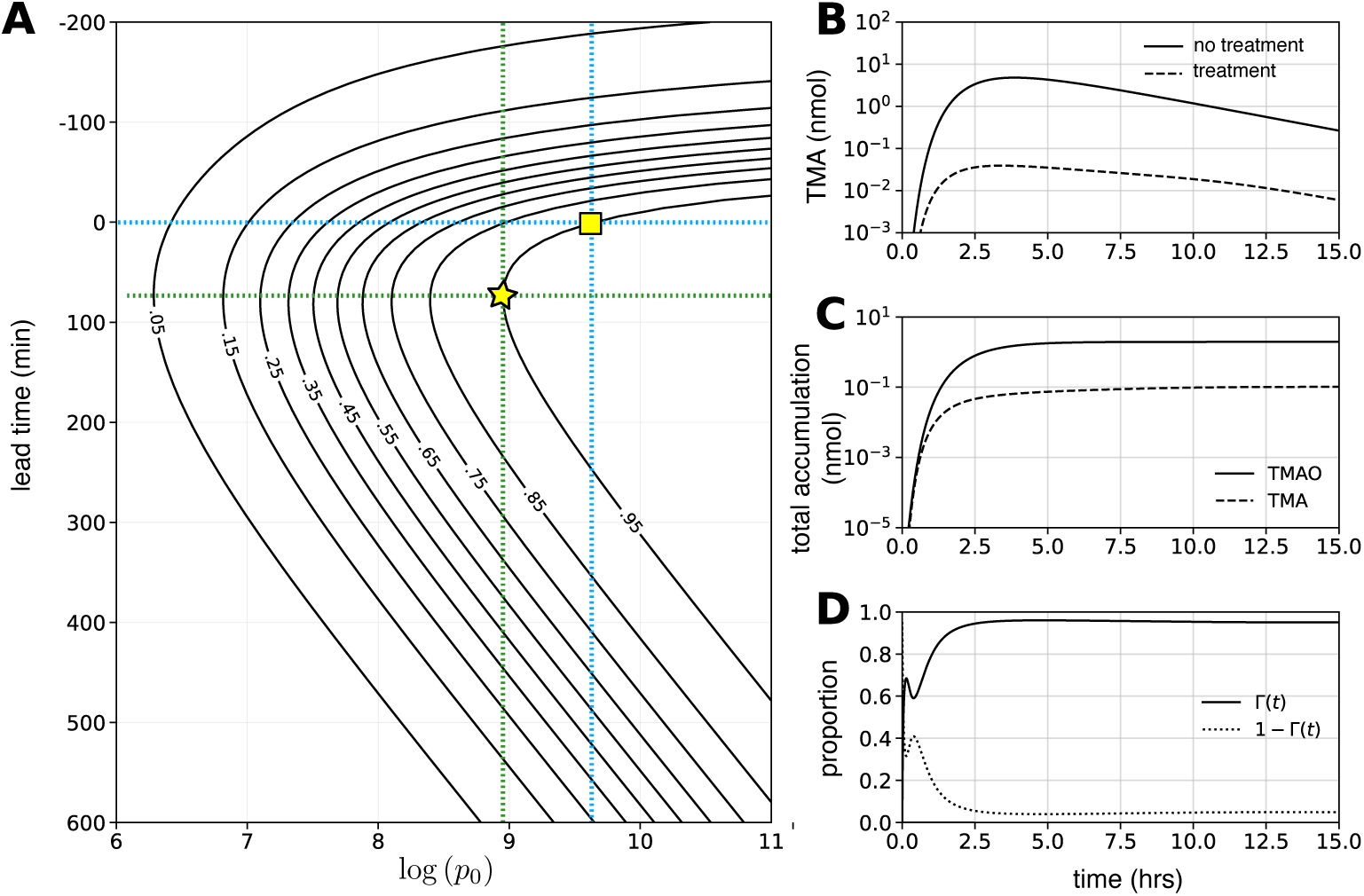
Treatment results. (A) Γ^∗^ values for a range of dose (*p*_0_) and lead time. Solid black lines are level curves of Γ^∗^ in *p*_0_ *−τ* parameter space. Yellow square indicates dosage required for healthy Γ^∗^ value if probiotic is ingested at same time as choline. Star indicates an optimum in the sense that the least amount of probiotic is required to obtain healthy levels of TMAO in the bloodstream. (B) Time series of TMA in the plasma in the treatment and no treatment scenarios. (C) Time series of total amount of TMA and TMAO accumulated in the plasma over time. (D) Time series of Γ(*t*) for *p*_0_ yielding Γ^∗^ = 0.95, indicating that convergence to 0.95 occurs within a few hours, indicating minimal relative TMA accumulation even in the transient.

The dose response is characterized in Figure 3A as a function of *p*_0_ and lead time (see ‘Earlier Probiotic Administration’). The dose response behavior for simultaneous probiotic and metabolite ingestion can be seen by tracking Figure 3A along the horiztonal line emanating from a lead time of 0. The key observation is that our model predicts that a probiotic dose of approximately 10^9.75^ CFUs will yield a 0.95 ratio of TMAO to TMA and TMAO in the plasma (yellow square). Though this number seems rather large, it is in complete agreement with other dosages for probiotics in other contexts [69]. For context, 10^9.75^ CFUs of probiotic can be integrated into a mixture that fits in a small capsule common in over the counter supplements. Maintenance of a Γ^∗^ value of 0.95 coincides with ratios observed in healthy individuals. Thus, the probiotic intervention met two goals: (1) it reduced TMA amount in the blood stream meaning fishy odor is diminished and (2) it allowed for the establishment of a Γ^∗^ of 0.95, signifying healthy metabolism.

#### 3.2.1 Earlier Probiotic Administration

The BCAT model is sufficiently flexible so as to also predict an optimal time for ingestion of probiotic. We demonstrate this here. In simulations described here, probiotic is administered at a time other than the exact time of choline (or more generally, metabolite) ingestion. If choline is ingested at time *T*, we administer probiotic at time *T* − *τ*, where we call *τ* the ‘lead time’. Here, *τ* ∈ R: *τ >* 0 indicates probiotic ingestion before metabolite consumption and *τ <* 0 indicates the opposite scenario.

In Figure 3A, we show dose response curves in the *p*_0_ − *τ* parameter space. Each solid line is a level curve parameterized by *p*_0_ and *τ* such that Γ^∗^ is constant. We generated these family of curves by systematically incrementing *τ* and *p*_0_ and solving the BCAT model till equilibrium is attained.

The BCAT model predicts that a lower dosage of probiotic is required for Γ^∗^ to equal 0.95 if it is administered at an earlier time. Specifically, our model predicts that a whole order of magnitude less of probiotic is required for healthy Γ^∗^ values to be obtained if probiotic is administered approximately 1 hr prior to choline consumption. This is significant because a key concern in biomolecular therapy and genetic engineering is the emerging drawbacks of probiotic ingestion. While generally speaking probiotics are encouraged among persons with gut issues, recent clinical trials have indicated that overconsumption of probiotics can result in increased susceptibility to antibiotic-resistant bacterial infection [70]. The scourge of antibiotic-resistant genes among microbial pathogens poses a serious threat to the effectiveness of current antimicrobial treatments, particularly for severe bacterial infections leading to sepsis [71]. Minimizing such risks is thus an important consideration in biomolecular therapy and genetic engineering.

The reason for earlier probiotic administration yielding the optimal outcome lies in the interplay of the kinetic parameters characterizing choline metabolism. *A priori*, gut bacteria are present in abundance throughout the small intestine, leading to the rapid metabolism of choline into TMA and subsequent flux into the plasma. Probiotic, on the other hand, must *establish* its presence throughout the small intestine, and while it flows through the small intestine at the same rate as choline (*K_t_*), the delay in access to TMA for oxidation stemming from metabolism of choline to TMA is just long enough to allow for more TMA flux into the plasma than is desired. Thus, simultaneous ingestion of choline and probiotic can suffice to reach healthy metrics, but it is necessary for high doses of probiotic, which, as we discussed above, may be undesirable.

On the other hand, earlier ingestion of probiotic allows it to establish its presence throughout the small intestine. Thus, as soon as TMA is available, probiotic can immediately begin oxidation, yielding desirable biometrics with smaller doses. Ingesting probiotic too early means the probiotic will be flushed out of the small intestine, causing a diminished presence at choline consumption. The optimal dosage occurs when these two opposing effects balance one another.

## 4 Conclusion

We developed a pharmacokinetic-pharmacodynamic hybrid model for probiotic intervention to treat metabolic disorders. Our model is based on the CAT model, which has been the source of successful studies in a pharmacokinetic capacity. In this paper, we provided a mathematical foundation for the origin of the CAT model, linking the successful compartment model to a PDE for linear transport. We further appended to the CAT model and incorporated bacterial metabolic impacts.

To our knowledge, our model is the first to mechanistically incorporate the metabolic effects of natural gut microbiota and to model probiotic transport and correlated metabolic effects upon biomolecular species under scrutiny. It is computationally cheap to simulate and, in particular instances, completely solvable. Our model is also general and flexible to incorporate a variety of enzymatic bioreactions. In our example, we used standard Michaelis-Menten kinetics, but using more sophisticated enzymatic metabolic pathways is straightforward in our framework. Moreover, in our model, we captured endogenous bacterial metabolism if choline with a mass action term. This primarily emphasizes that the relevant biophysical parameters pertaining to this metabolic pathway have not been reported. When such parameters are reported, incorporating correct enzymatic representations of the biocatalysis is also straightforward.

Here we demonstrated that the BCAT model can be used to ascertain two important quantities: (1) optimal dosage of an intervention to reach specific desired biometrics and (2) the optimal timing of the intervention. In particular, we applied the BCAT model to the specific metabolic disease TMAU with the intention of computing the dosage of synthetically engineering probiotic expressing TMM necessary for normal levels of TMA and TMAO to persist in the bloodstream. Our data-driven model and parameter selection predicts that ingesting approximately 1 billion CFUs of probiotic with a choline-rich meal suffices for the equilibrium TMAO/TMA ratio to reach the desired 0.95 quantity when probiotic is ingested approximately 1 hr before metabolite . While simultaneous administration of probiotic and metabolite requires higher dosages, the predicted amounts remain within practical and clinically achievable ranges. This important calculation will inform synthetic engineering required for design of TMM-expressing probiotic. Moreover, we believe our BCAT model can be used to inform dosages required for probiotic treatment of any metabolic disorder tied to gut microbiome defects.

Our model is relatively simple, and therefore there are a number of avenues to pursue in future work to increase the fidelity of our model to reality. We discussed how the kinetics of the various players (probiotic, choline) interplay to generate an optimal timing and dosage for desired biometrics. What we have not looked at here is the dependence of these optimal quantities with respect to the inherent system biophysical parameters. Which parameters are key for robust establishment of optimal intervention? Moreover, our model is quite coarse-grained. For example, we could incorporate diffusion of metabolite across the cell membrane of probiotic and model probiotic production of enzyme to more accurately capture the biophysical metabolic pathway. Nonetheless, our model can inform the design of the ultimate test of the accuracy of our model, which would be to actually synthesize the probiotic and test its efficacy. This was our original intention in undertaking this project. With a dosage calculation at hand, we will proceed in that direction.

## Acknowledgements

VLD is especially thankful for his parents, family and friends, who supported him throughout this work. BRK is eternally grateful for his wife, Hajra Habib, and his son, Surya.

## Footnotes

1 A CFU is a ‘colony-forming unit’ and is a measure of how many viable microbes are in a given population of microbes.

